# Quantitative fluorescence lifetime imaging uncovers a novel role for KCC2 chloride transport in dendritic microdomains

**DOI:** 10.1101/2020.06.08.139592

**Authors:** Nicholas L. Weilinger, Jeffrey M. LeDue, Kristopher T. Kahle, Brian A. MacVicar

**Affiliations:** Djavad Mowafaghian Centre for Brain Health, University of British Columbia, Vancouver, V6T 1Z3, Canada; Departments of Neurosurgery, Pediatrics, and Cellular & Molecular Physiology, Yale School of Medicine, Yale University, New Haven, CT, 06520, USA

**Keywords:** KCC2, Chloride, FLIM, NKCC1, MQAE, swelling, cytotoxic edema, dendrite, blebbing

## Abstract

Intracellular chloride ion ([Cl^−^]_i_) homeostasis is critical for synaptic neurotransmission yet variations in subcellular domains are poorly understood owing to difficulties in obtaining quantitative, high-resolution measurements of dendritic [Cl^−^]_i_. We combined whole-cell patch clamp electrophysiology with simultaneous fluorescence lifetime imaging (FLIM) of the Cl^−^ dye MQAE to quantitatively map dendritic Cl^−^ levels in normal or pathological conditions. FLIM-based [Cl^−^]_i_ estimates were corroborated by Rubi-GABA uncaging to measured E_GABA_. Low baseline [Cl^-^]_i_ in dendrites required Cl^−^ efflux via the K^+^-Cl^−^ cotransporter KCC2 (*SLC12A5*). In contrast, pathological NMDA application generated spatially heterogeneous subdomains of high [Cl^−^]_i_ that created dendritic blebs, a signature of ischemic stroke. These discrete regions of high [Cl^−^]_i_ were caused by reversed KCC2 transport. Therefore monitoring [Cl^−^]_i_ microdomains with a new high resolution FLIM-based technique identified novel roles for KCC2-dependent chloride transport to generate dendritic microdomains with implications for disease.

## Introduction

Neuronal function is intrinsically tuned by regulation of the intracellular Cl^−^ concentration ([Cl^−^]_i,_) which is critical for early brain development, setting membrane excitability, and cell volume regulation (Doyon et al., 2016; Kaila et al., 2014; Rungta et al., 2015). In mature nerve cells, low [Cl^−^]_i_ is maintained by K^+^-Cl^−^ cotransporter (KCC2)-dependent extrusion to set gamma aminobutyric acid (GABA)ergic inhibitory tone (Cordero-Erausquin et al., 2005; Kahle et al., 2013; Staley and Proctor, 1999). KCC2 is expressed throughout the dendritic arbor in hippocampal and cortical pyramidal neurons (Gauvain et al., 2011; Gulyas et al., 2001) and is highly expressed in synaptic regions in close proximity to N-methyl-D-aspartate (NMDA), GABA, and α-amino-3-hydroxy-5-methyl-4-isoxazolepropionic acid (AMPA) receptors to shape local excitatory/inhibitory potentials via shunting- and feeback-inhibition, and ionic plasticity (Chevy et al., 2015; Doyon et al., 2016; Garand et al., 2019; Gauvain et al., 2011). It is becoming clear that [Cl^−^]_i_ heterogeneity in neuronal subdomains is a key determinant of regional neurotransmission [(Barker and Ransom, 1978; Berglund et al., 2006; Glykys et al., 2014; Khirug et al., 2008; Kuner and Augustine, 2000), but also that local perturbations in [Cl^−^]_i_ may underlie synaptic dysfunction. For instance, KCC2 dysregulation has been implicated in multiple neurological disorders that involve synaptodendritic disinhibition, including epilepsy, neuropathic pain, schizophrenia, and autism (Cohen et al., 2002; Coull et al., 2003; Hyde et al., 2011; Steffensen et al., 2015; Tao et al., 2012). However, the spatial distribution and magnitude of dendritic [Cl^−^]_i_ changes in these diseases are poorly understood due to a lack of quantitative imaging of [Cl^−^]_i_ dynamics at the subcellular level.

The dynamic nature of Cl^−^ homeostasis makes it challenging to study using conventional electrophysiology or imaging techniques. Manipulating [Cl^−^]_i_ by whole-cell patch dialysis is difficult due to compensatory KCC2 or NKCC1 transport to re-establish physiological [Cl^−^]_i_ (Cordero-Erausquin et al., 2005; Doyon et al., 2016). Imaging genetically encoded fluorescent proteins like Clomeleons (Berglund et al., 2006; Grimley et al., 2013; Kuner and Augustine, 2000) and Cl^−^/H^+^ sensors (Mukhtarov et al., 2013; Sulis Sato et al., 2017) use ratiometric intensity emissions that are marred by divergent light scattering that is wavelength-, tissue depth-, and age-dependent (Oheim et al., 2001). These probes are also sensitive to pH (Arosio et al., 2007; Tsien, 1998), which is problematic given the interrelationship between Cl^−^ and HCO_3_^−^ cotransport (Kaila, 1994) and their joint permeabilities through many Cl^−^ channels (Bormann et al., 1987; Jun et al., 2016).

We adopted a Fluorescence Lifetime Imaging (FLIM)-based strategy to circumvent the limitations of ratiometric sensors and establish whether [Cl^−^]_i_ microdomains could be imaged. FLIM is insensitive to changes in dye concentration, light scattering, and photobleaching (Chen et al., 2013; Lloyd et al., 2010), making it ideal for quantifying [Cl^−^]_i_. The Cl^−^ sensor N-(ethoxycarbonylmethyl)-6-methoxyquinolinium bromide (MQAE) is suitable for FLIM with a broad dynamic range of lifetimes and is insensitive to pH (Gensch et al., 2015; Kaneko et al., 2002). Bulk-loaded MQAE has been used for quantitative Cl^−^ imaging *in situ* in Bergmann glia, dorsal root ganglion- and somatosensory neurons (Funk et al., 2008; Gilbert et al., 2007; Untiet et al., 2017), and in large dendritic knobs of the vomeronasal organ (Kaneko et al., 2004; Untiet et al., 2016). However, neurons bulk-loaded with MQAE have poor signal-to-noise over background staining, making it difficult to measure [Cl^−^]_i_ with synaptic resolution.

Here, we explored the utility of combining MQAE-FLIM with whole-cell patch clamp electrophysiology to measure dynamic variations in dendritic [Cl^−^]_i_ and the spatial distributions of these changes under both physiological and pathological conditions. We show MQAE-FLIM can accurately quantify [Cl^−^]_i_ subdomains in dendrites, and that basal [Cl^−^]_i_ is maintained even when experimentally challenged with elevated Cl^−^ loads. Using MQAE-FLIM, we quantified the relative contributions of KCC2 and NKCC1 in setting regional Cl^−^ electrochemical gradients, which we corroborated using Rubi-GABA uncaging to calculate E_GABA_ (Rial Verde et al., 2008). Lastly, we demonstrate that NMDA can generate localized dendritic microdomains with regions of remarkably high and persistent dendritic [Cl^−^]_i_ gradients. The formation of dendritic microdomains leads to the appearance of dendritic varicosities (“blebbing”) a hallmark of dendritic pathology. Remarkably, bleb formation was prevented by furosemide, indicating localized Cl^−^ loading was due to reversed K^+^/Cl^−^ transport. Together, our work highlights the advantages of MQAE-FLIM to measure [Cl^−^]_i_ shifts with sub-micrometer resolution that would be otherwise undetectable using intensity imaging or electrophysiology alone. Additionally, we have identified a novel role for KCC2 in dendritic microdomain Cl^−^ homeostasis that critically impacts neuronal structure and function.

## Results

### MQAE-FLIM calibrations for determination of [Cl^−^]_i_

As a first step we determined the reliability of MQAE-FLIM to report [Cl^−^]_i_ by calibrating the dye *in situ* (in bulk-loaded cells) and *in vitro* (in recording solution in sealed pipettes) by time-correlated single-photon counting using 2-photon microscopy (Fig 1a). *In situ* calibrations were conducted in acute brain slices bulk-loaded with MQAE using nigericin and tributyltin to equilibrate [Cl^−^]_i_ = [Cl^−^]_o_ across neuronal membranes (Gensch et al., 2015; Kovalchuk and Garaschuk, 2012). We then measured MQAE-FLIM lifetimes corresponding to various Cl^−^ concentrations (henceforth *MQAE-*[Cl^−^]_i_, Fig 1b,c) in parallel to intensity changes. Bulk-loaded MQAE intensity decreased significantly in response to increasing [Cl^−^]_o_ but we also observed progressive decline over time due to a combination of dye loss and photobleaching (Fig 1a,c). Indeed, intensity changes did not reverse in 0[Cl^−^]_o_ to wash out Cl^−^ (Fig 1d). In contrast, the MQAE-FLIM channel revealed discrete lifetime changes in response to increasing [Cl^−^]_o_ that fully recovered in 0[Cl^−^]_o_ washout despite the reduction in intensity signal (Fig 1d,e), highlighting the advantage and utility of FLIM in overcoming intensity artifacts arising from changes in dye concentration.

**Figure 1.**
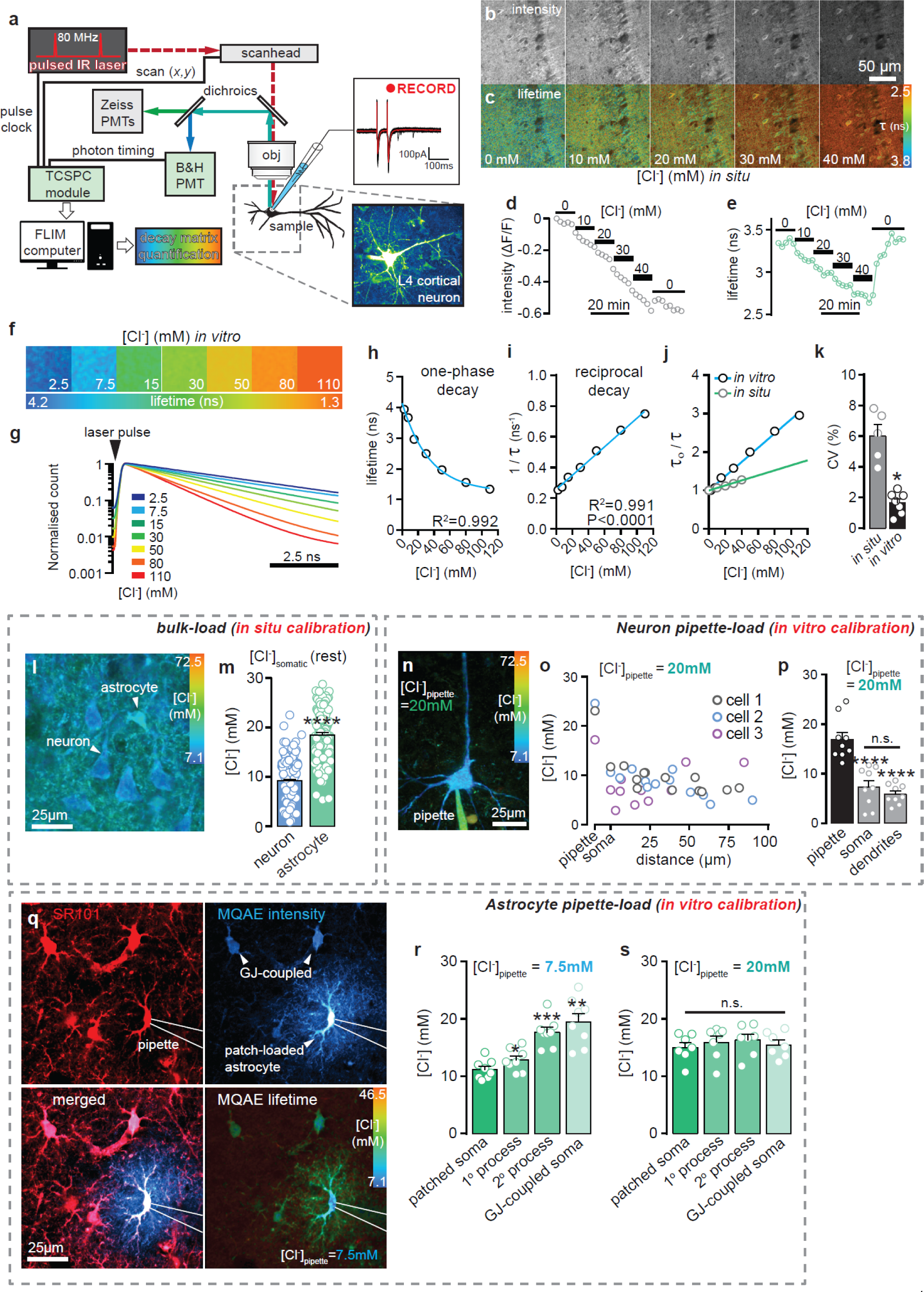
Fluorescence Lifetime Imaging reveals distinct Cl^−^ handling in neurons and astrocytes. **a)** Schematic of the time-correlated single photon counting FLIM setup pairing 2P laser imaging system with a Becker & Hickl SPC FLIM module and whole-cell patch clamp electrophysiology. **b)** MQAE-intensity image series of CA1 hippocampal neurons in the presence of nigericin and tributyltin to equilibrate [Cl^−^]_o_:[Cl^−^]_i_ across plasma membranes. [Cl^−^]_o_ was sequentially increased to alter [Cl^−^]_i_ and therefore MQAE-intensity. **c)** MQAE-FLIM readouts from images in ‘b’. **d)** Intensity measures from CA1 neuronal somata when [Cl^−^]_o_ is manipulated from 0 to 40 mM. Note that the signal does not reach steady state, nor does it recover to baseline upon washout. **e)** Corresponding MQAE-[Cl^−^]_i_ values for intensity measurements in ‘d’. Note that MQAE-FLIM signals reach steady state and that the Cl^−^ washout is quantifiable. **f)** Example FLIM images of *in vitro* MQAE calibration. **g)** Normalized MQAE lifetime curves at varying [Cl^−^]. **h)** MQAE-FLIM calibration data fit to a one-phase exponential decay curve consistent with a collisional quenching model. **i)** The reciprocal of the curve in ‘d’ confirms a linear relationship. **j)** KSV plots comparing calibrations obtained *in vitro* (in bulk-loaded cells) and *in vitro* (in sealed pipettes). **k)** Coefficient of variance (CV) plots comparing *in vitro* and *in situ* calibration data shows that the variability of lifetime readouts is lower *in vitro* (P<0.05, student’s t-test). **l)** Exemplar image of layer 4 cortical neurons and an astrocyte bulk loaded with MQAE. **m)** Mean somatic [Cl^−^]_i_ measurements with MQAE-FLIM reveal significantly higher basal [Cl^−^]_i_ in astrocytes (n=111 cells) compared to neurons (n=197 cells)(slices=6, ****P<0.0001, student’s t-test). **n)** Z-projection MQAE-FLIM image of layer 4 pyramidal neuron patch clamped with MQAE (6 mM) and set [Cl^−^]_i_ (20 mM) in the pipette. **o)** Quantification of [Cl^−^]_i_ plotted vs distance from the soma for three example cells to measure the precipitous drop in [Cl^−^]_i_ from the pipette to distal dendrites. **p)** Comparison of average [Cl^−^]_i_ readouts of all cells patched with [Cl^−^]_pipette_ = 20 mM (n=9 cells, ****P<0.0001, one-way ANOVA with Tukey test). **q)** Astrocytes identified by SR101 staining (top left) were patch clamped and dialysed with MQAE (top right) 7.5 mM Cl^−^. MQAE signal was detected in gap-junctionally (GJ) coupled astrocytes. **r)** Average MQAE-FLIM measurements comparing somatic [Cl^−^]_i_ to 1° and 2° processes (*P = 0.0171 and ***P = 0.0002, respectively), as well as to somatic ROIs from GJ-coupled astrocytes (**P = 0.0042) when [Cl^−^]_pipette_ = 7.5 mM (n= 8 slices, data compared to patched soma as a control, one-way ANOVA with Tukey test). **s)** Average MQAE-FLIM comparisons when [Cl^−^]_pipette_ = 20 mM (n= 7 slices, P>0.05, one-way ANOVA with Tukey test).

To combine MQAE-FLIM with whole-cell patch clamp, we conducted an additional *in vitro* calibration to complement and compare with the bulk-loaded *in situ* calibration. Whole cell loading provided improved signal to noise in dendritic compartments but it was important to calibrate MQAE in the electrode solution that included HEPES and gluconate (Kaneko et al., 2002). MQAE was dissolved in a K-gluconate recording solution with varied Cl^−^ concentrations, and MQAE-FLIM readouts were measured in sealed, temperature-controlled pipettes. Cl^−^-dependent MQAE lifetimes varied over a broad dynamic range, from ∼4.1 ns to ∼1.2 ns when [Cl^−^]_pipette_ was increased from 2.5 mM to 110 mM, respectively (Fig 1f-h). The calibration data were well-fit (R^2^ = 0.992) to a single exponential decay curve with a calculated *K*_*d*_ = 23.78 mM [Cl^−^] that should be suitable to capture physiological to pathological variations in [Cl^−^]_i_ (Fig 1h). The reciprocal of the dataset from Fig 1h correlated with a linear regression analysis (Fig 1i), confirming that MQAE works through collisional Cl^−^ quenching and therefore should not buffer Cl^−^ in solution (Verkman et al., 1989). As expected, the Cl^−^ sensitivity of MQAE differed in *in situ* and *in vitro* calibration environments (Stern-Volmer constants, K_SV_ = 6.53 M^-1^ and 20.19 M^-1^, respectively, Fig 1j), and the coefficient of variance was significantly lower in the *in vitro* setting (Fig 1k).

**Supplemental Figure 1.**
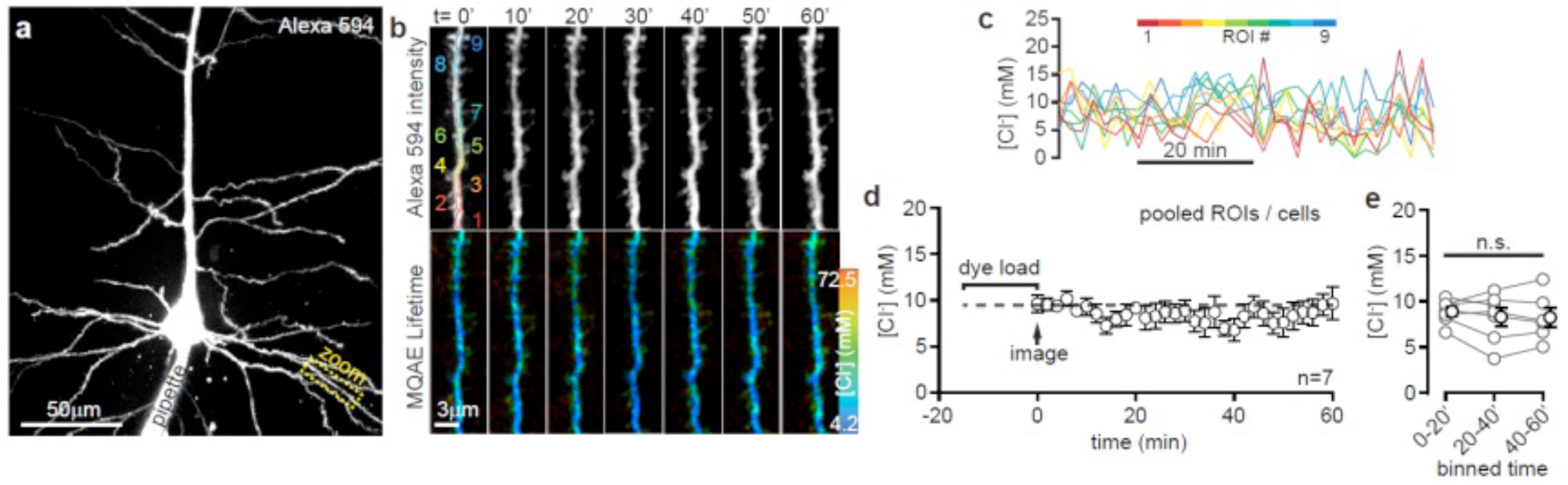
*MQAE-*[Cl^−^]_i_ readouts are stable in dendrites. **a)** Maximum intensity projection z-stack image of a cortical neuron patch-loaded with Alexa594 (presented emission) and MQAE. Dashed yellow box denotes basal dendrite imaged in ‘b’ and ‘c’. **b)** Sample Alexa 594 intensity (top panels) and MQAE-FLIM (bottom panels) image series to monitor dendritic morphology and maintain a stable focal plane for MQAE-FLIM. Intensity channel was used to map ROIs for MQAE-FLIM analysis. **c)** Color-coded MQAE-[Cl^−^]_i_ traces from dendrite in ‘b’. **d)** Average dendritic [Cl^−^]_i_ over time following a 15 min dye load period. Dendritic [Cl^−^]_i_ was stable over the course of 60 min imaging sessions (n=7 cells). **e)** Quantification of dendritic [Cl^−^]_i_ in 20 min intervals. No net change in [Cl^−^]_i_ was observed (P>0.05, Tukey’s test on one-way ANOVA), suggesting the relative impact of changes to the cytosolic microenvironment on MQAE-FLIM are negligible.

### Bulk- and patch-loaded cells report expected [Cl^−^]_i_ with MQAE-FLIM

Resting [Cl^−^]_i_ is reported to range from 6 mM to 13 mM in mature neurons (Delpire and Staley, 2014; Kuner and Augustine, 2000; Staley and Proctor, 1999). We sought to confirm that MQAE-FLIM measures of neuronal [Cl^−^]_i_ matched basal values reported in these previous studies. In neurons bulk loaded with MQAE, mean resting *MQAE-*[Cl^−^]_i_ was calculated to be 9.22±0.3 mM [Cl^−^]_i_ in the soma (perinuclear regions of interest [ROI]s) using *in situ* calibration values (Fig 1l,m). [Cl^−^]_i_ was significantly higher in astrocytes (18.48±0.5 mM) visually identified by cell morphology or by co-staining with SR101 (Fig 1l,m), closely matching measurements in cell culture (Gensch et al., 2015).

Next, we dialyzed MQAE by whole-cell patch clamp to compare MQAE-FLIM measures using *in vitro* calibration data in the soma and dendrites. Layer 4/5 cortical neurons were patch clamped with [Cl^−^]_pipette_ = 20 mM, a commonly used concentration with Ag/AgCl electrodes, and [Cl^−^]_i_ was measured at several distance intervals distal to the soma (Fig 1n,o). By calculating *MQAE-*[Cl^−^]_i_ with *in vitro* calibration values, we observed that somatic [Cl^−^]_i_ was consistently lower than the 20 mM [Cl^−^]_pipette_ and that the lower *MQAE-*[Cl^−^]_i_ readouts were stable throughout the proximal and distal dendrites (Fig 1n,o). Cellular [Cl^−^]_i_ in patched neurons closely matched measures from *in situ* bulk loading experiments (from Fig 1l,m), suggesting both methods accurately report resting [Cl^−^]_i_ levels (Fig 1p). As an important control, dendritic MQAE-FLIM readouts were stable over 1 hr (SFig 1), suggesting *MQAE-*[Cl^−^]_i_ measures are minimally affected by spatiotemporal variations of dye or dialysis of HEPES or gluconate from the pipette.

The precipitous drop from [Cl^−^]_pipette_ to somatic [Cl^−^]_i_ could be due to: (*1)* homeostatic Cl^−^ efflux by K^+^-Cl^−^ cotransport, or (*2)* unpredictable interactions between MQAE and the cytosolic microenvironment. The latter case assumes [Cl^−^]_i_ closely matches [Cl^−^]_pipette_ (20 mM) and that the measured *MQAE-*[Cl^−^]_i_ is artifactually altered by nonspecific anion quenching. We explored these possibilities by patch loading MQAE in astrocytes, which do not express KCC2 (Williams et al., 1999) and their [Cl^−^]_i_ is reported to range from 20-40 mM (Gensch et al., 2015; Untiet et al., 2017). Patched astrocytes were dialyzed with a [Cl^−^]_pipette_ concentration much lower than our *in situ* measure for astrocyte [Cl^−^]_i_ but closer to neuronal [Cl^−^]_i_ (7.5 mM). If hypothesis *(1)* was correct then homeostatic Cl^−^ import would increase [Cl^−^]_i_ to ∼18 mM (Fig 1l) due to high glial expression of NKCC1 (Su et al., 2002; Untiet et al., 2017). Somatic *MQAE-*[Cl^−^]_i_ in patched astrocytes was higher (11.16±0.6 mM) than [Cl^−^]_pipette_, consistent with the observation that astrocytes maintain a higher [Cl^−^]_i_ than neurons (Fig 1q,r)(Kettenmann and Schachner, 1985; MacVicar et al., 1989; Untiet et al., 2017). Distal measures revealed a progressive increase in [Cl^−^]_i_ in large primary-(1°) to smaller secondary (2°) processes of patched astrocytes, with peak measures from astrocyte somata coupled via gap-junctions (GJ)(19.46±1.46 mM, Fig 1r). In contrast, we observed a slight but reliable decrease in [Cl^−^]_i_ across all ROIs (Fig 1s) in astrocytes loaded with [Cl^−^]_pipette_ = 20 mM, consistent with an homeostatic decrease by KCC3 (Cordero-Erausquin et al., 2005; Doyon et al., 2016).

### Mapping subcellular KCC2 Cl^−^ export during intracellular Cl^−^ challenge

The extent of homeostatic Cl^−^ export via KCC2 was quantified using MQAE-FLIM by applying the inhibitor furosemide in neurons that were dialyzed with a high Cl^−^ load via the patch pipette ([Cl^−^]_pipette_ = 80 mM, normal aCSF [Cl^−^]_o_ = 136 mM). Under these conditions *MQAE-*[Cl^−^]_i_ was stable (somatic change = 2.74 ± 1.25 mM) after 15 min of dye loading (Fig 2ab), suggesting that MQAE diffusion was in stable equilibrium and that its lifetime is not appreciably impacted by hydrolysis (Koncz and Daugirdas, 1994). We again observed a marked decrease in somatic *MQAE-*[Cl^−^]_i_ compared to *MQAE-*[Cl^−^]_pipette_ (Fig 3ac) with gradual, distance-dependent reductions in dendrites suggesting substantial Cl^−^ efflux throughout the dendritic tree. Bath application of furosemide (200 µM) significantly increased dendritic [Cl^−^]_i_ at binned ROIs equidistant to the soma (Fig 2c-e), indicating that the low dendritic [Cl^−^]_i_ required KCC2. Applying furosemide together with picrotoxin to inhibit GABA_A_-receptors similarly increased [Cl^−^]_i_ by blocking tonic Cl^−^ efflux from GABA_A_R when *E*_*Cl-*_ is depolarized from holding potential (*V*_*m*_= -60 mV) (Fig 2d,e). Interestingly, [Cl^−^]_i_ was significantly lower in 2° versus 1° dendrites (Fig 2f), consistent with the observation that KCC2 is evenly distributed in dendrites and therefore Cl^−^ handling should be more efficient where the surface area to volume ratio is high (Doyon et al., 2011). These data suggest neurons defend their Cl^−^ gradients by KCC2-transport.

**Figure 2.**
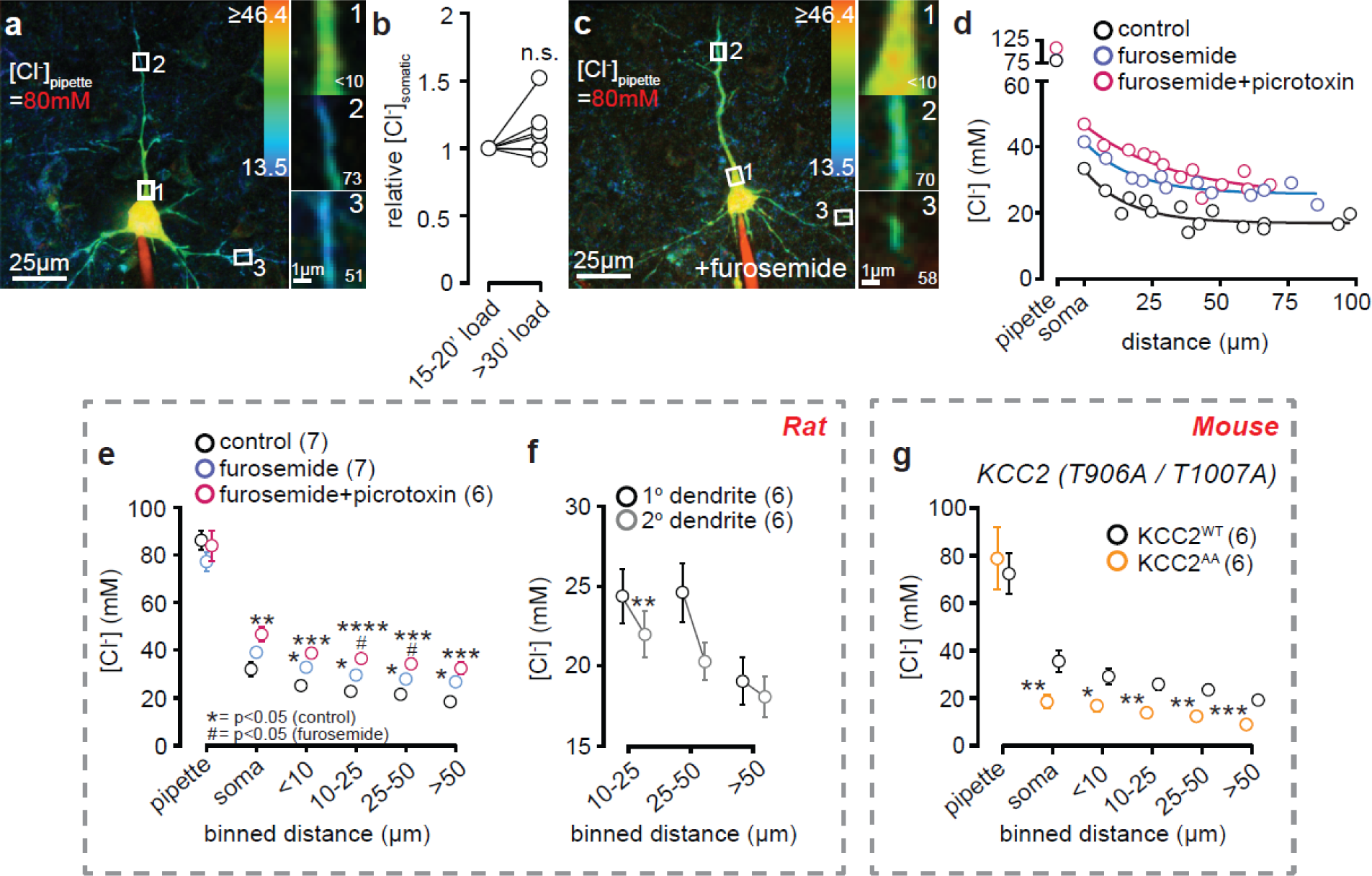
KCC2-dependent Cl^−^ extrusion during high [Cl^−^]_i_ challenge. **a)** Left panel: example maximum intensity projection (collapsed Z-stack) image of layer 4 neuron patch filled with [Cl^−^]_pipette_ = 80 mM, normal ASF [Cl^−^]_o_ = 136 mM, voltage clamped at -60 mV. Right panels: zoomed ROIs from primary (panels 1&2) and secondary (panel 3) dendrites at different distances from the soma (in µm, bottom right values in figure panels). **b)** Quantification of average [Cl^−^]_i_ measures over time (each point represents whole-cell averages). [Cl^−^]_i_ stabilized after 15 min post whole-cell break-in, and MQAE-FLIM readouts did not change 30 min post break-in. **c)** Exemplar cell loaded with [Cl^−^]_pipette_ = 80 mM in the presence of KCC2 blocker furosemide (200 µM), normal ASF [Cl^−^]_o_ = 136 mM, voltage clamped at -60 mV. Zoom panels show higher steady-state [Cl^−^]_i_ measured by MQAE-FLIM compared to ‘a’, suggesting a KCC2-dependent Cl^−^ efflux pathway. **d)** Plots of [Cl^−^]_i_ at varying distances from the soma. Application of furosemide alone, or furosemide+picrotoxin (100 µM) increased steady state [Cl^−^]_i_ measures by MQAE-FLIM compared to control. **e)** Quantitative summary comparing drug treatments on [Cl^−^]_i_ at respective distances (*P<0.05 compared to control, ^#^P<0.05 compared to furosemide). **f)** Comparison of [Cl^−^]_i_ between primary and secondary dendrites at binned ROIs equidistant from the soma. [Cl^−^]_i_ measures were relatively lower in 2° vs 1° dendrites at distances of 10-25 µm (**P = 0.0062) from the soma, but not at distances 25-50 µm (P = 0.0517) and >50 µm (P = 0.2467), two-tailed paired t-tests. **g)** Quantitative analysis of [Cl^−^]_i_ by MQAE-FLIM in KCC2^A/A^ mice compared to WT controls patch loaded with [Cl^−^]_pipette_ = 80 mM, normal ASF [Cl^−^]_o_ = 136 mM, voltage clamped at -60 mV. [Cl^−^]_i_ was significantly lower in KCC2^A/A^ neurons compared to KCC2^WT^ (from left to right, P = 0.0082, 0.0161, 0.0014, 0.0012, 0.0006), consistent with an increased extrusion capacity of KCC2.

**Figure 3.**
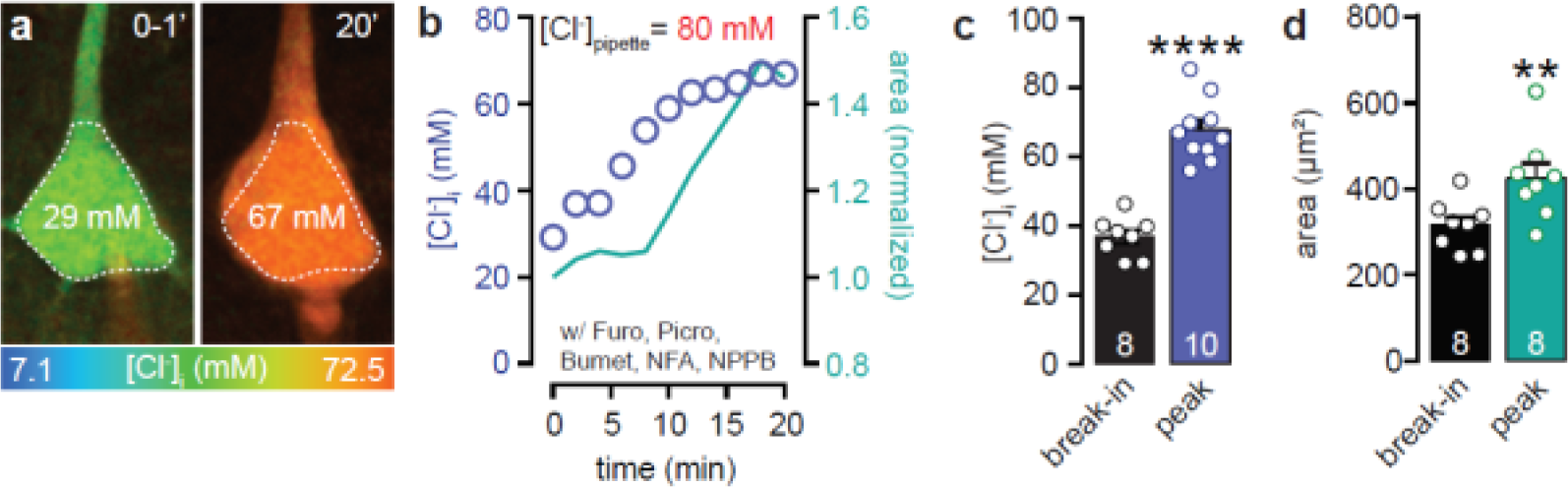
Inhibiting putative routes of Cl^−^ efflux leads to dramatic increase in [Cl^−^]_i_. **a)** Example MQAE-FLIM images at whole-cell break in (left panel) and 20 min after (right panel) of cell patch clamped with [Cl^−^]_pipette_ = 80 mM, normal ASF [Cl^−^]_o_ = 136 mM, voltage clamped at -14 mV. Experiment performed with furosemide (200 µM), picrotoxin (100 µM) bumetanide (50 µM), NFA (100 µM), and NPPB (100 µM) in the ACSF. **b)** Quantification of somatic [Cl^−^]_i_ and cross-sectional area over time in ‘a’. **c)** Summary of peak [Cl^−^]_i_ measures by MQAE compared to whole-cell break-in (0-1 min)(****P<0.0001, two-tailed paired t-test). **d)** Comparison of somatic swelling as measured by cross-sectional area (**P = 0.0018, two-tailed paired t-test).

KCC2 transport activity is highly regulated by kinase-dependent phosphorylation via the With No-Lysine (K) (WNK) and Ste20p-related Proline Alanine-rich Kinases (SPAK) (Alessi et al., 2014). Phosphorylation or dephosphorylation of threonines (T) T906 and T1007 on the KCC2 c-terminus cause transporter hypo- or hyperfunction, respectively (Friedel et al., 2015; Kahle et al., 2013; Rinehart et al., 2009). To further explore KCC2’s role in establishing Cl^−^ gradients, we repeated the above experiments in knock-in mice in which T906 and T1007 were mutated to alanine (A) (i.e., T906A and T1007A, henceforth “KCC2^A/A^”) to mimic KCC2 dephosphorylation and constitutive activation (Friedel et al., 2015; Heubl et al., 2017; Moore et al., 2018; Pisella et al., 2019; Watanabe et al., 2019). We observed a sharp drop in somato-dendritic *MQAE-*[Cl^−^]_i_ in patched KCC2^A/A^ neurons compared to wild-type controls (KCC2^WT^), consistent with an increased pump capacity of KCC2^A/A^ (Fig 2g). Additionally, we found no difference in equidistant [Cl^−^]_i_ measures between pyramidal neurons in brain slices from control rat versus mouse (KCC2^WT^) (control and KCC2^WT^ groups from Fig 2e,g, respectively, P>0.05 by two-way ANOVA). This similarity suggests that the KCC2 extrusion capacity is consistent between these species. We conclude that MQAE-FLIM can detect dynamic fluctuations in [Cl^−^]_i_ through manipulation of KCC2 activity, and that discrepancy between *MQAE-*[Cl^−^]_i_ and *MQAE-* [Cl^−^]_pipette_ is primarily due to homeostatic Cl^−^ extrusion.

If the *MQAE-*[Cl^−^]_pipette_:*MQAE-*[Cl^−^]_i_ mismatch was due to the cytosolic environment, somatic [Cl^−^]_i_ readouts from Fig 2 would be closer to 80 mM instead of the recorded ∼30-40 mM, and true [Cl^−^]_i_ would be close to a ceiling and likely not amenable to increase. We tested whether *MQAE-*[Cl^−^]_i_ in these conditions could be increased to match [Cl^−^]_pipette_ by experimentally inhibiting Cl^−^ fluxes. Neurons were voltage clamped at the calculated *E*_*Cl-*_ for [Cl^−^]_pipette_ = 80 mM (*E*_*Cl-*_ = -14 mV), and a cocktail of blockers was applied to inhibit Cl^−^ fluxes, including: picrotoxin (GABA_A_R), furosemide, bumetanide (NKCC1), niflumic acid (NFA, Ca^2+^-activated Cl^−^ channels), and 5-Nitro-2-(3-phenylpropylamino)benzoic acid (NPPB, VRACs). Under these conditions, somatic [Cl^−^]_i_ steadily increased from 36.83±2.1 mM at break-in to a plateau of 64.11±1.8 mM over 20 min (Fig 3a-c). This was accompanied by a volume increase (as evidenced by the increase in mean cross sectional area = 34.8%, Fig 3a,b,d) because Cl^−^ loading is a key driver of swelling (Rothman, 1985; Rungta et al., 2015). Together, these data refute hypothesis *(2)* and suggest that *MQAE-*[Cl^−^]_i_ readouts can be used to quantify Cl^−^dynamics via transporter export and efflux through Cl^−^ channels.

### Local GABA uncaging corroborates MQAE lifetime

As final confirmation that MQAE-FLIM is accurate, we compared spatially coupled FLIM readouts to E_GABA_ measured by local two-photon of uncaging Rubi-GABA (Rial Verde et al., 2008). Rubi-GABA (0.5 mM) was locally applied and photolyzed adjacent to the somatic MQAE-FLIM ROI. In this way we would achieve reasonable voltage clamp in the soma with minimal space clamp artifacts (Fig 4a) (Williams and Mitchell, 2008). These experiments were performed in the absence of the intracellular K^+^-channel blocker Cs^+^ to prevent perturbation of KCC2 function (Blaesse et al., 2009; Williams and Payne, 2004). In neurons patched with [Cl^−^]_pipette_= 80 mM, Rubi-GABA was uncaged at -80 mV to +20 mV and [Cl^−^]_i_ was calculated from *E*_*GABA*_ (*E*_*GABA*_-[Cl^−^]_i_, Fig 4b-d). *E*_*GABA*_-[Cl^−^]_i_ and *MQAE-*[Cl^−^]_i_ values were well correlated (Fig 4f) despite known confounds for *E*_*GABA*_ measures such as permeability to HCO_3_^−^ (Kaila and Voipio, 1987; Kaila et al., 1993). To test the impact Br^−^ efflux (dissociated from MQAE-Br salt) on *E*_*GABA*_, we reduced the concentration of MQAE 10-fold and found no statistical difference from 6 mM MQAE (Fig 4e).

**Figure 4.**
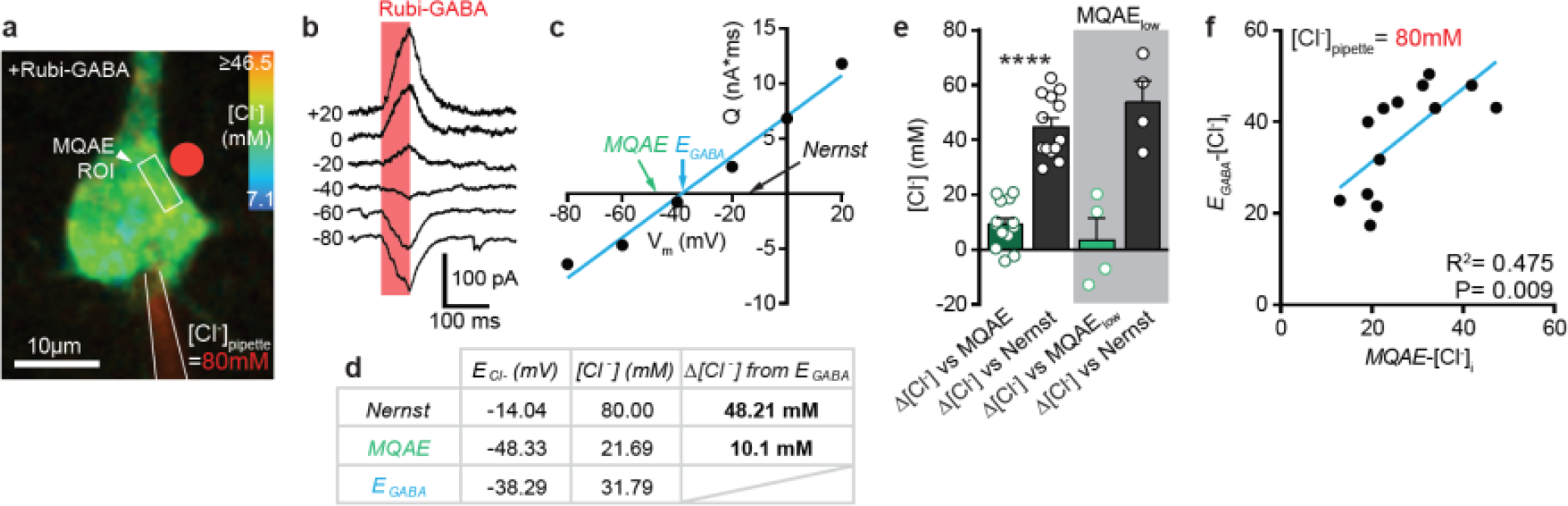
Comparison of MQAE-[Cl^−^]_i_ to E_GABA_-[Cl^−^]_i_. **a)** Example image of layer 4 neuron patch-loaded with [Cl^−^]_pipette_ = 80 mM, normal ASF [Cl^−^]_o_ = 136 mM. Rubi-GABA was locally applied (0.5 mM) by puff electrode and photolyzed adjacent to the soma (red dot) by 2P-uncaging. **b)** Example whole-cell currents induced by Rubi-GABA uncaging (red box) at different holding potentials. **c)** Charge (Q) and membrane voltage (V_m_) plot of Rubi-GABA responses. EGABA is graphically compared to Cl^−^ reversal calculated by MQAE measurement, as well the E_Cl_-estimation that assumes [Cl^−^]_i_ = [Cl^−^]_pipette_ (80 mM). **d)** Tabular results from exemplar experiment ‘a-c’ showing the calculated values for E_Cl_-based on the pipette solution (Nernst), the MQAE-FLIM measurement, and the E_GABA_ measurement; the respective predicted [Cl^−^]_i_ values; and the differences in the predicted [Cl^−^]_i_. **e)** Comparison of [Cl^−^]_i_ predicted by E_GABA_ to *MQAE*-[Cl^−^]_i_ and Nernst, ****P<0.0001. **f)** Correlation plot between calculated *E*_*GABA*_-[Cl^−^]_i_ and *MQAE*-[Cl^−^]_i_ values.

### Mapping KCC2/NKCC1-dependent control of dendritic Cl^−^ gradients

We measured the basal contributions of NKCC1 and KCC2 in setting dendritic [Cl^−^]_i_. Proximal basal dendrites ([Cl^−^]_pipette_ = 7.5 mM) were imaged by MQAE-FLIM before and after bath application of furosemide or the NKCC1 inhibitor bumetanide (Fig 5a-d). Quantitative and reversible increases in *MQAE-*[Cl^−^]_i_ were observed in furosemide, consistent with disruption of basal Cl^−^ efflux by KCC2 (Fig 5a-c). On the other hand, bath application of bumetanide (50 µM) to selectively block NKCC1 decreased resting [Cl^−^]_i_ due to reduced Cl^−^ import. As a positive control, membrane depolarization to +30 mV also increased [Cl^−^]_i_ in the presence of furosemide or bumetanide (Fig 5c,d) but also in the absence of either blocker (Fig 5e), consistent with an increased driving force for Cl^−^ entry through tonic GABA_A_R activity and voltage-activated Cl^−^ channels. Thus, MQAE-FLIM has the sensitivity to measure millimolar contributions of Cl^−^ channels and transporters, as well as dynamic [Cl^−^]_i_ changes.

**Figure 5.**
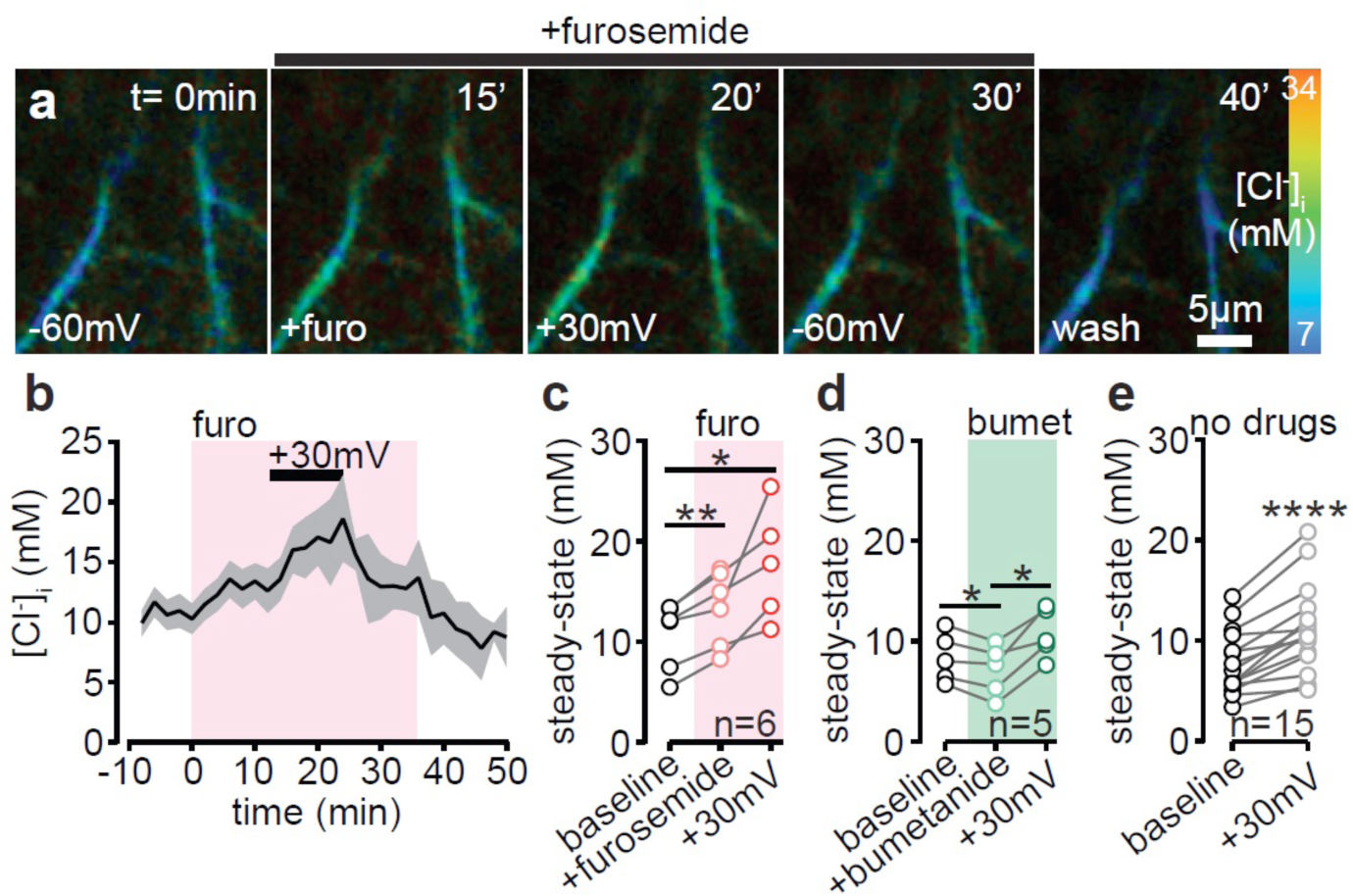
MQAE-FLIM can detect KCC2-dependent changes in [Cl^−^]_i_ with millimolar precision. **a)** Example MQAE-FLIM image series of [Cl^−^]_i_ in a basal dendrite from a patch clamped layer 4 cortical neuron ([Cl^−^]_pipette_ = 7.5 mM) in the presence of furosemide (furo, 200 µM) and membrane depolarization (+30 mV). **b)** Average trace of [Cl^−^]_i_ changes over time in response to furosemide application (pink box) and membrane depolarization (n=6). **c)** Quantitative analysis comparing paired [Cl^−^]_i_ measurements before and after furosemide (**P = 0.006) and membrane depolarization to +30 mV (*P = 0.0378), one-way ANOVA with Tukey test. **d)** Quantitative comparison of paired [Cl^−^]_i_ measurements before and after bumetanide (bumet, 50 µM, *P = 0.026) and membrane depolarization to +30 mV (*P = 0.0165), one-way ANOVA with Tukey test. **e)** Membrane depolarization alone is sufficient to increase dendritic [Cl^−^]_i_, ****P<0.0001, two tailed paired t-test.

### MQAE-FLIM reveals heterogeneous Cl^−^ microdomains in cytotoxic edema

The importance of Cl^−^ loading in cytotoxic swelling is well established (Rothman, 1985; Rungta et al., 2015) although little is known about the spatiotemporal dynamics and regulation of subcellular Cl^−^ loading in dendritic regions during edema. Excitotoxic injury in ischemia is thought to be triggered by NMDA receptor overstimulation leading to dendritic swelling and blebbing (Murphy et al., 2008; Thompson et al., 2008; Weilinger et al., 2016; Weilinger et al., 2012). We used MQAE-FLIM to monitor [Cl^−^]_i_ dynamics under intense NMDA receptor stimulation in dendrites to recapitulate this form of ischemic injury. We used a strategy that we previously developed to restrict the actions of bath applied NMDA to only single patched clamped neuron (Dissing-Olesen et al., 2014). In these experiments 6 mM Mg^2+^ was included in the aCSF perfusate to block tissue-wide NMDAR responses from bath applied NMDA. Only single L4/5 cortical neurons that were patch clamped and depolarized to +30 mV to relieve the NMDAR Mg^2+^ block responded to NMDA application (20 µM) (Dissing-Olesen et al., 2014). In this way, swelling was restricted to the patched neuron throughout the experimental time course, thereby providing the stability to simultaneous track *MQAE-*[Cl^−^]_i_ and morphological changes in dendritic regions over time in the intact brain slice.

NMDA induced a rapid increase in *MQAE-*[Cl^−^]_i_ followed by characteristic dendritic blebbing (Fig 6a,b) in depolarized neurons. We observed a surprising, heterogeneous distribution of [Cl^−^]_i_ in excitotoxic conditions, with higher [Cl^−^]_i_ constrained to dendrites that formed blebs (Fig 6c-e). Indeed, [Cl^−^]_i_ in blebs was significantly higher than adjacent dendrites that did not bleb, often reaching peak concentrations of ∼80 mM (Fig 6d,e). Pixel-to-pixel lifetime measurements showed that Cl^−^ microdomains were spatially restricted across the length of dendrites (Fig 6f,g). We resolved precipitous drops in [Cl^−^]_i_ spanning a measured bleb edge length of <1 µm (mean [Cl^−^]_i_ decrease = -26.32±7.9 mM/µm)(Fig 6h), suggesting that the mobility of Cl^−^ ions is constrained during blebbing.

**Figure 6.**
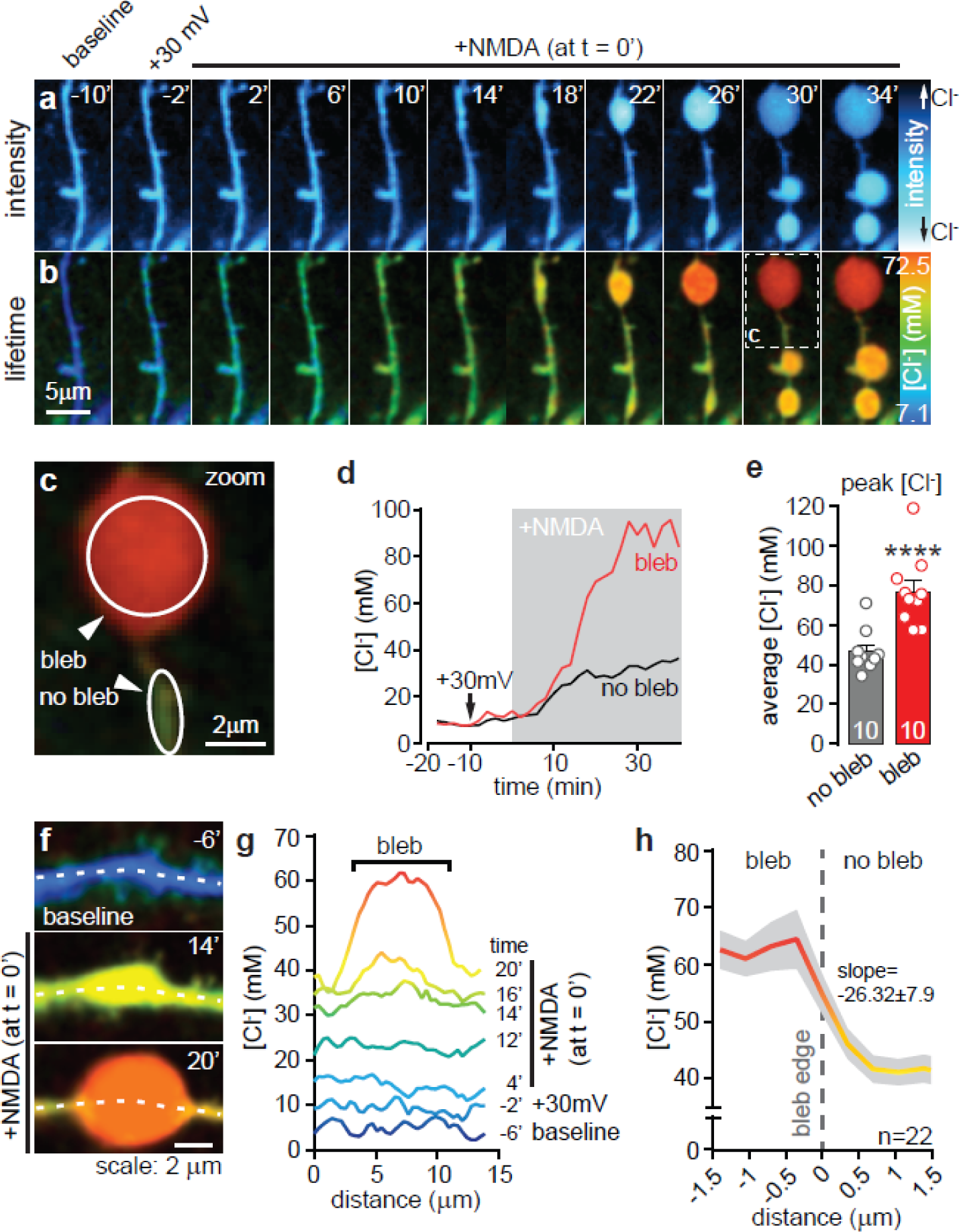
MQAE-FLIM reveals heterogeneous [Cl^−^]_i_ landscapes in excitotoxicity. **a)** Example imaging time series of MQAE intensity in dendrite from a patch-clamped neuron ([Cl^−^]_pipette_ = 7.5 mM) before and after membrane depolarization and bath application of NMDA (20 µM). **b)** MQAE-FLIM image data (same dendrite from ‘a’) reveals dramatic [Cl^−^]_i_ heterogeneities during dendritic blebbing that are not resolved from intensity images. **c)** Zoomed image of dendrite from ‘b’ (white box t= 30 min) depicting example ROIs encompassing blebbed and adjacent un-blebbed dendrite. **d)** Representative [Cl^−^]_i_ measurements over time from bleb and no bleb ROIs in the presence of NMDA. **e)** Quantitative summary of peak [Cl^−^]_i_ measures comparing MQAE-FLIM signal from blebbed and unblebbed ROIs, ****P<0.0001. **f)** MQAE-FLIM time course showing the formation of a bleb. Time (in min) in the top right corner of the images indicates the time relative to application of NMDA (at t = 0 min). Dotted white line indicates the ROI used to measure the pixel-to-pixel [Cl^−^]_i_ levels over distance reported in ‘g’. **g)** Spatial [Cl^−^]_i_ readouts from before, during, and after formation of the bleb shown in ‘f’. Each line represents a [Cl^−^]_i_ measurement along a dendrite in a given frame. Corresponding time points are shown to the right of each trace. NMDA is applied at t = 0 min. **h)** Average ± s.e.m. of MQAE-FLIM [Cl^−^]_i_ readouts across bleb edges reveal a sharp decrease [Cl^−^]_i_ in from bleb to no bleb that occurs over less than a 1 µm span. ‘n’ denotes 22 blebs from 9 cells.

### Excitotoxic Cl^−^ dysregulation in dendrites is KCC2-dependent

The mechanistic underpinnings of Cl^−^ entry in dendritic blebbing is a point of contention. Previous work has identified roles for VRACs in cultured neurons *in vitro*, and KCCs and GABA_A_ receptors in acute brain slices (Allen et al., 2004; Inoue and Okada, 2007; Pond et al., 2006; Steffensen et al., 2015). We sought to dissect which Cl^−^ transporter or channel activity engendered the abnormally high but discrete [Cl^−^]_i_ accumulations in dendritic blebs. We first confirmed the critical role for Cl^−^ entry in swelling by reducing extracellular Cl^−^ from 136.5 mM to 18.5 mM (low[Cl^-^]_o_). Low[Cl^-^]_o_ blocked the Cl^−^ influx as well as blebbing (Fig 7a-d) down to control levels (+30 mV only group, Fig 7d). Swelling also requires Na^+^-influx to drive Cl^−^ entry and satisfy Gibbs-Donnan equilibrium (Glykys et al., 2017). Consistent with this, reducing Na^+^ in otherwise normal ACSF decreased NMDA-induced Cl^−^ influx (Fig 7a-d) and blebbing, confirming that concurrent Na^+^ and Cl^−^ loading is required.

**Figure 7.**
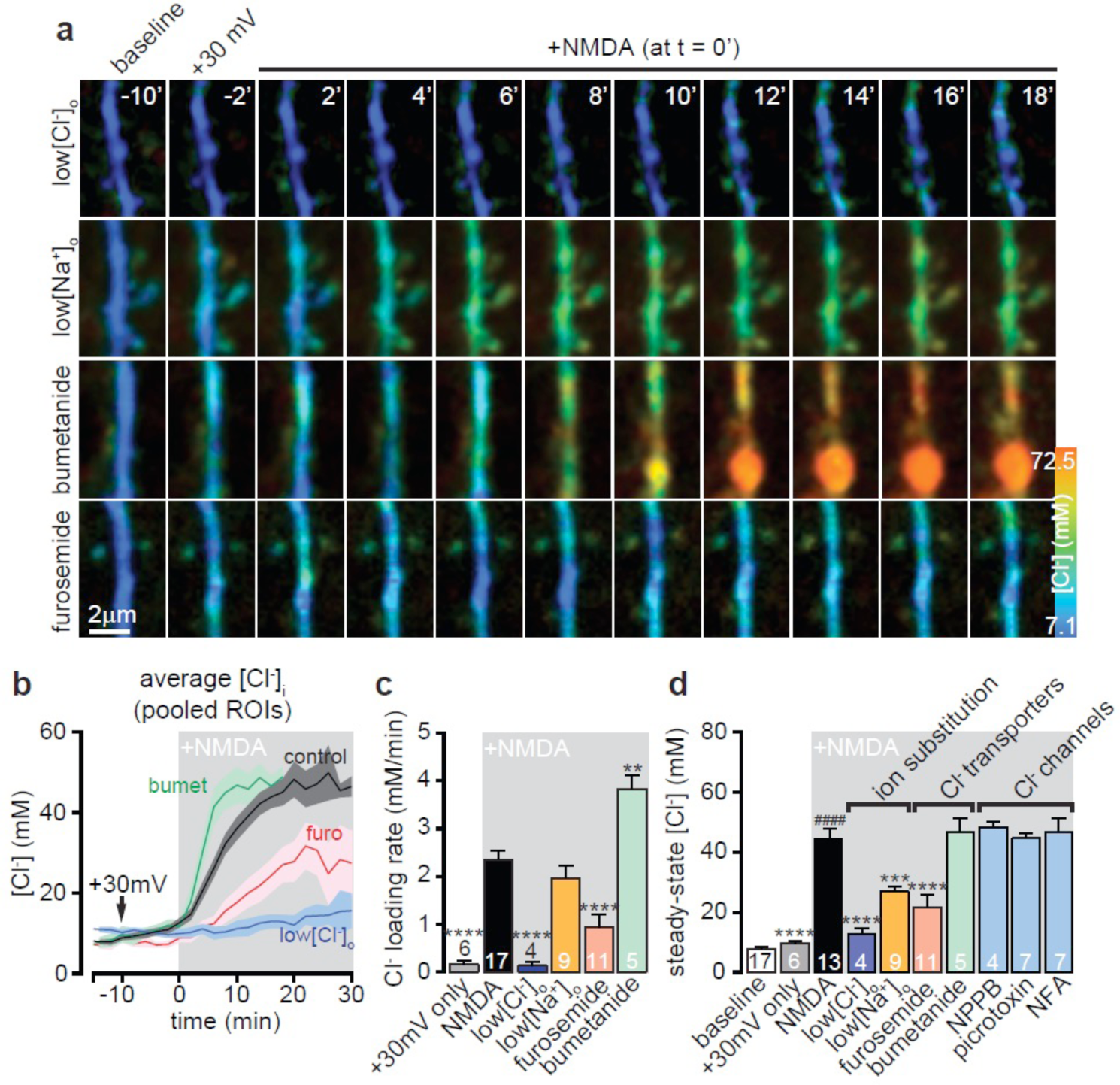
NMDA-induced Cl^−^ influx and blebbing is KCC2 dependent. **a)** Example MQAE-FLIM time-lapses of NMDA-induced blebbing in low[Cl^−^]_o_, low[Na^+^]_o_, bumetanide, and furosemide conditions. NMDA (20 µM) was continually applied at time = 0 min. **b)** Average *MQAE-*[Cl^−^]_i_ readouts over time for datasets outlined in ‘a’. Data are pooled ROI averages (i.e. encompassing blebbed and non-blebbed ROIs). **c)** Average Cl^−^ loading rate, calculated from the rising slope of NMDA-induced Cl^−^ increase from ‘b’. +30 mV only indicates continuous depolarization for 40 min. ****P<0.0001 for NMDA alone compared to +30mV, [Cl^−^]_o_, and furosemide conditions, **P = 0.0032 compared to bumetanide, one-way ANOVA with Tukey test. **d)** Quantitative summary of steady-state *MQAE-*[Cl^−^]_i_ at 15 min post NMDA application. ^####^P<0.0001 NMDA alone vs baseline, ****P<0.0001 significance vs NMDA alone, one-way ANOVA with Tukey test.

We tested the role for NKCC1 in mediating Cl^−^ entry in swelling by pre-incubating bumetanide prior to NMDA application. Interestingly, bumetanide increased the rate of Cl^−^ loading during the initial NMDA application but the peak [Cl^−^]_i_ accumulations during the plateau were not different from NMDA alone (Fig 7b,c). In contrast, blocking KCC2 with furosemide dramatically reduced the Cl^−^ loading rate and average [Cl^−^]_i_ load (Fig 7a-d). Cl^−^ channel inhibitors, such as 5-Nitro-2-(3-phenylpropylamino)benzoic acid (NPPB, 100 µM) to block VRACs, picrotoxin (100 µM) for GABA_A_ receptors, and niflumic acid (NFA, 100 µM) to block Ca^2+^ activated Cl^−^ channels had no effect on NMDA-induced Cl^−^ loading. Together, these data highlight the importance of the local regulation of Cl-microdomains to maintain dendrite volume. Further, our findings suggest that Cl^−^ dysregulation in excitotoxic blebbing occurs principally through reversed uptake by KCC2.

### Legends for movies 1, 2 and 3 that provide time lapse illustrations for data in Figure 7

**Movie 1: NMDA triggers dendritic [Cl–]i loading and blebbing**. Time lapse of MQAE-FLIM images to measure spatiotemporal changes in dendritic [Cl–]i. NMDA (20 µm) application occurs at ‘+NMDA’ tag and is immediately followed by dramatic and localized increases in [Cl–]_i_ that form dendritic blebs.

**Movie 2: Dendritic blebbing requires Cl– influx**. Application of NMDA (20 µm) does not generate substantial subcellular increases in low [Cl–]o conditions and dendritic blebbing does not occur, indicating that dendritic NMDA-induced blebbing requires Cl– influx.

**Movie 3: Blocking KCC2 with furosemide reduces NMDA-induced Cl– influx and blebbing**. Bath pre-application of furosemide (200 µM) dramatically reduced Cl– loading and blebbing in the presence of NMDA (20 µm), indicating that KCC2 transport direction is reversed in cytotoxic edema.

## Discussion

Early observations of subcellular [Cl^−^]_i_ heterogeneities using electrophysiology (Alger and Nicoll, 1979; Barker and Ransom, 1978; Nicoll et al., 1976) have been corroborated by recent studies employing ratiometric sensors (Berglund et al., 2006; Glykys et al., 2014; Kuner and Augustine, 2000; Sulis Sato et al., 2017), however our understanding of the spatial regulation and functional impact of these microdomains is limited. Here we demonstrate that pairing MQAE-FLIM with electrophysiology allowed us to provide the first high resolution quantitative measurements of the spatiotemporal [Cl^−^]_i_ dynamics throughout cortical pyramidal dendrites. MQAE-FLIM had both the sensitivity and dynamic range to measure equilibrative [Cl^−^]_i_ changes under resting conditions and cytotoxic changes during intense NMDA receptor stimulation. We quantified the basal contributions of NKCC1 and KCC2 in maintaining [Cl^−^]_i_ at rest as well as modest changes in dendritic [Cl^−^]_i_ incurred by membrane depolarization. Importantly, we report that NMDA-induced cytotoxic swelling is driven by Cl^−^ influx via KCC2 reversed transport. Our observations lead to the surprising conclusion that discrete microdomains of high [Cl^−^]_i_ can be generated with sharp boundaries that give rise to dendritic blebbing.

Quantifying dendritic Cl^−^ dynamics and spatial distributions have been longstanding challenges. This is due to the robust homeostatic control of Cl^−^ gradients, as well as the bidirectional nature of Cl^−^ flux that depends on dynamic *E*_*m*_ and *E*_*Cl–*_ values (Arosio and Ratto, 2014). Additionally Cl^−^ imaging using many genetically encoded fluorescent proteins is problematic due to pH sensitivity, but also non-uniform distortions caused by the optical properties of brain tissue that alter even ratiometric excitation and emission values (Oheim et al., 2001; Sulis Sato et al., 2017). This is further complicated by volume changes that occur in parallel with [Cl^−^]_i_, which can affect intrinsic optical properties of the tissue and thereby subsequent intensity changes. Two recent studies show substantial improvements in this area using dual ratiometric Cl^−^-pH sensors such as ClopHensorN and LSSmClopHesnsor (Mukhtarov et al., 2013; Sulis Sato et al., 2017). However MQAE-FLIM is still preferable for quantitative studies due to its high Cl^−^ affinity, fast detection kinetics, insensitivity to changes in emission intensity, dye concentration, photobleaching, and light scattering (Arosio and Ratto, 2014; Fukuda et al., 1998; Kovalchuk and Garaschuk, 2012; Marandi et al., 2002; Verkman et al., 1989). MQAE is also relatively insensitive to pH fluctuations (Koncz and Daugirdas, 1994) and viscosity (Kaneko et al., 2002). In support of this, we observed a near-complete inhibition of NMDA-induced rises in *MQAE*-[Cl^−^]_i_ in low[Cl^−^]_o_, even though NMDA triggers cytosolic acidification (Canzoniero et al., 1996). One limitation for imaging MQAE is nonspecific but modest quenching by gluconate, EGTA and HEPES (but notably not by endogenous anions such as SO_4_^2–^, HCO_3_^−^, and PO_4_^3–^) (Kaneko et al., 2002; Kovalchuk and Garaschuk, 2012). This is typically controlled for by calibrating the dye *in situ*, although complete [Cl^−^]_i_:[Cl^−^]_o_ equilibration is difficult to achieve. Indeed, our measured K_SV_ for *in situ* (6.53 M^-1^) and *in vitro* (20.19 M^-1^) calibrations were both within the range of *in situ* experiments from another group (5-20 M^-1^)(Kaneko et al., 2004) and well below MQAE in distilled water (185 M^-1^). It is therefore beneficial to patch load MQAE, as these unspecific anion interactions can be controlled for by their inclusion in calibration solutions.

Notwithstanding the challenges and considerations associated with lifetime imaging, our MQAE-FLIM readouts from bulk- and pipette-loaded configurations are in agreement with each other and with basal [Cl^−^]_i_ values predicted in the literature (Delpire and Staley, 2014; Kuner and Augustine, 2000; Staley and Proctor, 1999). An important validation of our approach is its accuracy compared to regional *E*_*GABA*_ measures. *MQAE*-[Cl^−^]_i_ readouts were <10 mM below *E*_*GABA*_-[Cl^−^]_i_, which we attribute to a combination of GABA_A_R HCO_3_^−^ efflux (Kaila, 1994) and imperfect voltage clamp due to K^+^ leak in the absence of K^+^ channel blockers. Nevertheless, the variance in both our *MQAE*-[Cl^−^]_i_ and *E*_*GABA*_-[Cl^−^]_i_ values were strongly correlated, and commensurate with the physiological heterogeneity in [Cl^−^]_i_ reported by other groups (Barker and Ransom, 1978; Delpire and Staley, 2014; Glykys et al., 2014; Khirug et al., 2008; Kuner and Augustine, 2000; Sulis Sato et al., 2017). There was also no detectable impact of Br^−^ ion flux either *MQAE*-[Cl^−^]_i_, meaning MQAE-Br could be used over a broad concentration range depending on experimental requirements. The stability of *MQAE*-[Cl^−^]_i_ over a 60 min recording indicates that quenching from endogenous and exogenously (pipette) supplied anion species was minimal relative to Cl^−^. Thus, the variability of [Cl^−^]_i_ we observe within neuronal dendrites highlights the clear need for quantitative evaluations of the functional implications of Cl^−^ microdomains.

The temporal dynamics of Cl^−^ handling has been shown to be modified over time scales of days (during development), minutes (KCC2 posttranslational regulation by calpains or WNK/SPAK), and milliseconds-seconds (ionic plasticity, shunting) (Alessi et al., 2014; Doyon et al., 2016; Kaila et al., 2014). With MQAE-FLIM we quantified transport capacity of NKCC1 and KCC2 over time but also at multiple subcellular sites. For example, patched neurons and astrocytes defended their respective low (<10 mM) and high (15-20 mM) [Cl^−^]_i_ setpoints when dialyzed with mismatched Cl^−^ loads. Interestingly, we observed regional variance in Cl^−^ export in both cell types, with higher transport efficiency in dendrites and fine processes. MQAE-FLIM also enabled us to measure a wide range of [Cl^−^]_i_ changes. By blocking KCC2 or NKCC1 in dendrites with either furosemide or bumetanide, we could quantify subtle (<5 mM) changes in Cl^−^ transport at rest with basal synaptic transmission intact. Detection of such modest [Cl^−^]_I_ variations is testament to the sensitivity and practicability of MQAE-FLIM, as millimolar changes would affect excitability (Doyon et al., 2016). We also measured large [Cl^−^]_i_ changes in neurons loaded with [Cl^−^]_pipette_ = 80 mM, where somatic and dendritic was reduced typically below 40 mM in part by export via KCC2. This allowed us to also quantify the impact of KCC2 dephosphorylation in T906A/T1007A mice (Friedel et al., 2015; Heubl et al., 2017; Moore et al., 2018; Pisella et al., 2019; Watanabe et al., 2019), where [Cl^−^]_i_ was maintained at near-control levels despite an elevated [Cl^−^]_pipette_ load (80 mM). Indeed, there is growing appreciation for dynamic KCC2 phosphorylation, for example in establishing excitatory GABAergic tone during development, impacting cognitive function, and susceptibility to epileptiform activity (Friedel et al., 2015; Moore et al., 2019; Moore et al., 2018; Pisella et al., 2019; Silayeva et al., 2015; Watanabe et al., 2019).

Evidence of spatial [Cl^−^]_i_ heterogeneity within cellular subcompartments in previous studies (Barker and Ransom, 1978; Berglund et al., 2006; Glykys et al., 2014; Khirug et al., 2008; Kuner and Augustine, 2000) suggested that standing Cl^−^ gradients could exist within cytoplasmic domains of neurons. Our observations of inhomogeneous [Cl^−^]_i_ distributions in dendrites at rest and the dramatic induction of discrete [Cl^−^]_i_ microdomains that lead to dendritic blebs with distinct edges indicates that Cl^−^ mobility can be spatially restricted. Although controversial (Kaila et al., 2014), the suggestion that Cl^−^ ions are diffusion-limited, even at rest, has been proposed to occur due to a nonuniform distribution of immobile intracellular and extracellular anions [(Glykys et al., 2014) see also co-submission by Rahmati et al., 2020]. While our data cannot confirm the role of immobile anions in setting Cl^−^ microdomains, the uneven distribution cytosolic proteins (Gut et al., 2018) could give rise to compensatory heterogeneities in mobile Cl^−^ ions. In cytotoxic edema, breakdown of negatively charged proteins and structural actin is well established in excitotoxicity (Halpain et al., 1998; Lipton, 1999) and would likely affect Cl^−^ mobility in dendrites (Glykys et al., 2014). The possibility of Cl^−^ microdomains might be expected in some dendritic regions given that dendritic morphology alone can affect ion diffusion. For instance, the thin necks of synaptic spines can restrict Ca^2+^ diffusion and propagation of electrical potentials between spine heads and dendrites (Kwon et al., 2017; Yuste and Denk, 1995). Additionally, recent modelling studies predict that longitudinal Cl^−^ diffusion in dendrites is restricted by the distribution and strength of K^+^-Cl^−^ cotransporters (Doyon et al., 2011), by tortuosity imposed spines (Mohapatra et al., 2016), and even by passive membrane properties (Lombardi et al., 2019), all of which could affect local GABAergic inhibition and ionic plasticity. Given that little is known about Cl^−^ microdomains and their regulation at the sub-dendritic level, future studies using MQAE-FLIM will be fundamental to understanding KCC2’s vital role in synaptic physiology.

The data presented here indicate that formation of pathological blebs requires reversed KCC2 transport that generates microdomains of high dendritic [Cl^−^]_i_. Quantitative readouts of dendritic [Cl^−^]_i_ revealed spatially discrete increases in Cl^−^ that were blocked by furosemide, leading to our conclusion that KCC2 drives Cl^−^ import into blebs. We also observed a surprising increase in Cl^−^ loading rate when NKCC1 was blocked with bumetanide, indicating that NKCC1 was also pumping in reverse mode (i.e. Cl^−^ export) which has been suggested to occur when Na^+^, K^+^, and Cl^−^ concentration gradients are dramatically altered (Brumback and Staley, 2008; Glykys et al., 2017; Kaila et al., 2014). KCC2 transport direction depends on the outwardly-directed K+ gradient but also the stoichiometry of K^+^:Cl^−^ ions (Payne, 1997). *In silico* and *in vitro* experiments on KCC2 thermodynamic driving force have shown K^+^/Cl^−^ cotransport can reverse when [K^+^]_o_ increases beyond 5 mM (DeFazio et al., 2000; Payne, 1997). Reversed transport is therefore likely to occur in excitoxicity/stroke, where [K^+^]_o_ increases to >60 mM (Hansen and Nedergaard, 1988; Rossi et al., 2007). A recent study showed that brief ischemia induced prolonged (>1hr) increases in neuronal [Cl^−^]_i_ and epileptiform activity that were sensitive to furosemide block (Blauwblomme et al., 2018). Further, the frequently observed decrease in KCC2 expression in neurotraumas (Coull et al., 2003; Jaenisch et al., 2010; Kaila et al., 2014; Zhou et al., 2012) could be explained by the present findings as a protective adaptation to minimize pathological Cl^−^ loading in neurons. In conclusion, our joint MQAE-FLIM and electrophysiology approach has provided valuable insights into the tight spatiotemporal regulation of [Cl^−^]_i_ microdomains as it relates to integration of synaptic currents, but also in neuropathologies where perturbed Cl^−^ handling can degrade inhibitory tone in epilepsy or trigger dendritic swelling in cerebral edema (Cohen et al., 2002; Rungta et al., 2015).

## Methods

### Animals

All animal care protocols were approved by the University of British Columbia’s Animal Care Committee in accordance with the Canadian Council on Animal Care guidelines. Acute slice experiments from rat were performed on postnatal day (P) >25 Sprague Dawley rats (Charles River). Mouse experiments were performed on P>25 KCC2^AA^ (homozygous) or KCC2^WT^ animals. All animals were housed on a 12/12hr light/dark cycle with at least one cage mate and *ad libidum* access to laboratory chow and water.

### Acute Cortical Slice Preparation

Rats were anaesthetized by isoflourane inhalation in air and decapitated. The brain was quickly extracted, blocked, and mounted on a vibrating slicer (Leica VT1200S) while submerged in an ice-cold solution consisting of (in mM): 120 NMDG, 2.5 KCl, 25 NaHCO_3_, 1 CaCl_2_, 7 MgCl_2_, 1.25 NaH_2_PO_4_, 20 glucose, 2.4 Na-pyruvate, 1.3 Na-ascorbate and saturated with 95% O_2_/5% CO_2_. Transverse cortical slices were cut (370 µm) and transferred into a chamber containing artificial cerebral spinal fluid (aCSF) at 33°C for at least one hour prior to experimentation. aCSF consisted of 126 mM NaCl, 26 mM NaHCO_3_, 2.5 mM KCl, 1.25 mM NaH_2_PO_4_, 2 mM MgCl_2_, 2 mM CaCl_2_, and 10 mM glucose and was continuously bubbled with 95% O_2_/5% CO_2_.

### Chemicals and Reagents

Drugs and dyes used for experiments are listed in final concentration (in mM) and were purchased from the following suppliers: 0.2 furosemide, 0.1 niflumic acid (Cayman Chemicals), 0.05 bumetanide, 0.05 picrotoxin, 0.02 NMDA, 0.01 tributyltin, all salts for aCSF and slicing solution (Sigma-Aldrich), 0.01 Glycine (EMD Chemicals), 0.1 NPPB, 0.05 bumetanide, and 0.01 nigericin (Tocris), 0.05 Alexa-594 hydrazide, 0.6-8 MQAE (Thermo Fisher Scientific). All drugs were made into aliquots dissolved in water or DMSO and diluted to the final concentration in aCSF or intracellular recording solution – the final concentration of DMSO never exceeded 0.1%.

### Electrophysiology

After stabilization, slices were carefully transferred to a recording chamber continually perfused with aCSF (33°C, stage heater from Luigs & Neumann) at a rate of 1-2 mL/min. Layer 4/5 pyramidal neurons in the cortex were identified using widefield infrared (IR) illumination and captured with an IR-1000 (DAGE-MTI) camera fitted on a LSM 7MP 2-photon imaging system (Zeiss). Whole-cell patch clamp recordings were obtained using thin-walled borosilicate glass microelectrodes (Warner) pulled to a tip resistance of 3-5 MΩ pulled using a P-97 Flaming/Brown Micropipette Puller (Sutter Instrument). Electrodes were filled with an intracellular recording solution containing (in mM): 108 K-Gluconate, 3 KCl, 2 MgCl_2_, 8 Na-Gluconate, 1 K_2_-EGTA, 0.23 CaCl_2_, 0.05 Alexa-594, 6-8 MQAE, 4 K_2_-ATP and 0.3 Na_3_-GTP at pH 7.25 with 10 HEPES. 15 minutes was allotted for intracellular equilibration for all experiments unless otherwise indicated. Recordings were made using a MultiClamp 700B amplifier and Digidata 1440A digitiser (Axon Instruments, Molecular Devices) controlled via Clampex 10.7 acquisition software. Cells were voltage clamped at -60 mV unless specified otherwise. Access resistance was always <20 MΩ, and cells with holding currents below -100 pA after break-in were discarded.

### Time-Resolved 2-photon fluorescent lifetime imaging setup

All experiments were performed on a LSM 7MP 2-photon imaging system from Zeiss fitted with a SPC-150 FLIM acquisition module from Becker & Hickl. Femtosecond excitation was achieved with a Ti:Sapphire Chameleon Ultra II 2-photon laser (Coherent) pulsing at 80MHz and tuned to 750nm. Laser pulse timing was collected directly from the laser’s internal photodiode. Images were acquired with a Zeiss 20X-W/1.0 NA objective at digital zooms ranging from 3x-12x at 256×256 or 128×128 pixel resolutions depending on zoom factor. Emission light was first passed through an IR filter (700nm shortpass) to block spurious excitation light, then split with a 480nm longpass dichroic mirror (Chroma tech). Blue MQAE fluorescence was then filtered with a 460/50nm bandpass filter and collected with a GaAsP hybrid detector (HPM-100-40 hybrid PMT, Becker and Hickl). After the 480nm beam splitter, red emission from Alexa-594 was passed through a 630/75nm bandpass filter (Chroma) before detection with a LSM BiG GaAsP photomultiplier (Zeiss).

### MQAE-FLIM Calibrations

*In situ* calibrations were conducted as previously described (Gensch et al., 2015; Kovalchuk and Garaschuk, 2012) to measure [Cl^−^]_i_ in neurons and astrocytes bulk loaded with MQAE. Briefly, acute transverse hippocampal/cortical slices were incubated in 6 mM MQAE for 20 min at 33-34°C. Slices were then transferred to the imaging chamber containing the Cl^−^/OH^−^ antiporter tributyltin and the K^+^/H^+^ ionophore nigericin in a 0-40 mM KCl solution also containing 10 mM HEPES, 10 mM Na-Gluconate, pH balanced to 7.35 with KOH, osmolarity adjusted to 300 with K-Gluconate, and warmed to 33°C.

*In vitro* MQAE calibrations were performed to match the recording conditions in whole-cell patch clamp experiments. Calibration solutions exactly mimicked the internal recording solution (see ‘*Electrophysiology*’) including Alexa-594. The desired Cl^−^ concentration was achieved by adjusting the concentration of KCl and was balanced by equimolar K-gluconate. For 2.5mM [Cl^−^], some MgCl_2_ was substituted with MgSO_4_, as MQAE fluorescence is not quenched by SO4^2–^ anions (Kaneko et al., 2002). The calibration solution was placed in a sealed micropipette and imaged in warmed (33°C) aCSF under the microscope using consistent experimental parameters as *in situ*. Calibration data were fit in GraphPad Prism 6 with a one-phase exponential decay curve to match the monoexponential decay parameters of MQAE lifetime:

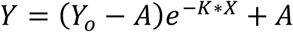

Where Y = measured lifetime (in ps), X = [Cl^−^], Y_o_ = lifetime at 0[Cl^−^] (in ps), A = lifetime horizontal asymptote at high [Cl^−^] (in ps), K = rate constant. Our best-fit line calculated from pooled averages was:

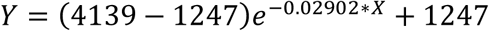

Due to the limitations of the curve fit, any measured lifetime values that were above the upper limit of 4139 ps were set to 0 mM [Cl^−^]. The *K*_*d*_ value for Cl^−^-dependent MQAE quenching was calculated as *K*_*d*_=0.69/K.

### Fluorescent Lifetime Imaging and Data Analysis

For bulk loaded imaging, slices were loaded with MQAE (6 mM) as described above. Neuronal somata were identified visually, and astrocytes were identified by SR101 stain (1 µM, post-slice incubation) or visually in unstained tissue. Images were acquired by continual XY frame scanning over 15s-60s to ensure adequate photon counts for accurate lifetime curve-fitting at low laser power. All cells were imaged between 50-100µm below the slice surface to balance minimal fluorescent scattering and maximal cell viability. For patch loaded imaging, cells were dialyzed with internal solution containing 6 mM MQAE for a least 15 min prior to imaging to allow for complete equilibration of dye and pipette salts.

Lifetime data were collected in parallel to signal intensity for each pixel and analysed, and the intensity data were used to process lifetime data and for illustration purposes. Lifetime data were processed using SPCImage 7.3 software from Becker & Hickl and decay values of each pixel were calculated based on a monoexponential fit described above. A bin factor of 1 was used to attain a photon count ≥10 at the tail (9ns after laser pulse) and a χ^2^ fit value close to 1. Lifetime decay matrices were decimated from 32 bit to 16 bit by rounding values to the nearest 1 ps in MATLAB (Mathworks). Intensity images were used as a registration template to align the lifetime images in MATLAB, and dendritic lifetime signals were measured in ImageJ. For dendritic recordings, ROIs were mapped on intensity images to ensure signal measurements were in-focus and transferred to the lifetime image. All colour-coded example FLIM images in the figures are intensity-weighted RGB images for ease of visualisation of MQAE signal over background fluorescence.

### Rubi-GABA uncaging

Rubi-GABA (Rial Verde et al., 2008) was purchased from Tocris and dissolved directly into recording aCSF at 0.5 mM and stored as frozen aliquots prior to use. The Rubi-GABA solution was always protected from ambient light, during preparation and experimentation. Rubi-GABA was locally puff-ejected near the patched neuron (∼100 µm) and held at a constant pressure of 1 psi throughout the experiment. Somatic MQAE-FLIM measures were then taken with the 2-photon laser tuned to 750 nm and imaged at low power (<4 mW) to prevent GABA uncaging during the acquisition. The laser was then tuned to 800 nm for Rubi-GABA photolysis, and uncaging laser power was slowly increased until reliable GABA_A_ receptor currents were observed at *V*_*m*_ = -80 mV (50 ms pulses). Patched neurons were filled with 8 mM MQAE in 80 mM Cl^−^ intracellular recording solution, and 0.5 µM TTX was included in normal aCSF to block voltage-gated Na^+^ channels. A liquid junction potential of 6 mV was applied to all voltage steps. *E*_*GABA*_-[Cl^−^]_i_ was calculated from *E*_*GABA*_ using the Nernst Equation.

### Dendritic blebbing assay

To isolate swelling in a single cell we used a high Mg^2+^ (6 mM) aCSF solution to increase the likelihood of voltage-dependent Mg^2+^ block of tissue-wide NMDA receptors as previously described (Dissing-Olesen et al., 2014). Layer 4/5 cortical neurons were whole-cell patch clamped with MQAE in a K-gluconate internal solution described above and held at *V*_*m*_ = -60 mV. Cells were dialyzed for a 10-20 min baseline period prior to imaging, followed by a 10 min depolarization to *V*_*m*_ = +30 mV to ensure complete unblock NMDA receptors in dendrites and accommodate loss of membrane potential control from space clamp issues. Sustained bath application of 20 µM NMDA and 10 µM glycine was used to trigger Cl^−^ loading and swelling. Low [Cl^−^]_o_ aCSF was made my replacing NaCl with equimolar Na-Gluconate, and low [Na^+^]_o_ aCSF by replacing NaCl with NMDG^+^ and pH balanced with HCl. Apical dendrites were selected for imaging based on their proximity to the soma for voltage clamp (<100 µm), and images were acquired over 20-30s, typically 160 frames, every 2 min to minimize phototoxicity.

## Acknowledgements

N.L.W collected and analyzed the data and wrote and edited the manuscript with B.A.M. N.L.W and K.K. designed the study with B.A.M., and B.A.M. supervised the study.

